# The presence or absence alone of miRNA isoforms (isomiRs) successfully discriminate amongst the 32 TCGA cancer types

**DOI:** 10.1101/082685

**Authors:** Aristeidis G. Telonis, Rogan Magee, Phillipe Loher, Inna Chervoneva, Eric Londin, Isidore Rigoutsos

## Abstract

Previously, we demonstrated that miRNA isoforms (isomiRs) are constitutive and their expression profiles depend on tissue, tissue state, and disease subtype. We have now extended our isomiR studies to The Cancer Genome Atlas (TCGA) repository. Specifically, we studied whether isomiR profiles can distinguish amongst the 32 cancers. We analyzed 10,271 datasets from 32 cancers and found 7,466 isomiRs from 807 miRNA hairpin-arms to be expressed above threshold. Using the top 20% most abundant isomiRs, we built a classifier that relied on “binary” isomiR profiles: isomiRs were simply represented as ‘present’ or ‘absent’ and, unlike previous methods, all knowledge about their expression levels was ignored. The classifier could label tumor samples with an average sensitivity of 93% and a False Discovery Rate of 3%. Notably, its ability to classify well persisted even when we reduced the set of used features (=isomiRs) by a factor of 10. A counterintuitive finding of our analysis is that the isomiRs and miRNA loci with the highest ability to classify tumors are <u>not</u> the ones that have been attracting the most research attention in the miRNA field. Our results provide a framework in which to study cancer-type-specific isomiRs and explore their potential uses as cancer biomarkers

## INTRODUCTION

In the post-genomic era, the flourishing of microarray and next-generation sequencing (NGS) technologies has made the generation of large amounts of data a quick and relative inexpensive process. As a result, the challenge nowadays is how best to manipulate and analyze large volumes of information. This has led to the development of novel tools, or the adaptation of previously-developed algorithms for use in the biological and medical fields (1).

Throughout this era, the field of non-coding RNAs (ncRNAs) enjoyed very significant progress. In fact, RNA-sequencing technologies helped uncovered many novel categories of short and long RNA transcripts (2). In the process, they also revealed multiple layers of regulatory processes.

Among ncRNAs, microRNAs (miRNAs) are arguably the best studied to date (3–5). The details of miRNA biogenesis (6–8) and function (7,9,10) were worked out more than a decade ago. Parallel studies linked miRNAs to a wide range of cellular, molecular and physiological processes in development (11–15), and homeostasis (16–18). In addition, miRNAs play important roles in physiological conditions and diseases (19–21), including cancer (22,23).

Their potent regulatory roles, small size, and relatively easy quantification have made miRNAs ideal targets as potential biomarkers (24–26). MiRNAs have also inspired research on their use as tumor classifiers. i.e. as features/variables used to construct statistical models to classify and/or predict the type of a given tumor. For example, Lu *et al.* used hierarchical clustering to classify miRNA profiles of tumor samples into groups of cancer types (27). Subsequent work by Volinia *et al*. (28) and by Rosenfeld *et al*. (29) further demonstrated the power of the miRNA profile to classify tumors and predict cancer types.

More than two decades since their discovery, the mining of RNA-seq datasets continues to generate important observations about miRNAs. Perhaps most important is the ever increasing repertoire of miRNAs, which has implications for the complexity of the miRNA regulatory layer. This was recently demonstrated by the discovery of numerous primate-specific miRNAs with tissue-dependent expression patterns (30). This complexity increased further with the recent discovery that miRNA isoforms (isomiRs) are constitutive and that isomiR expression depends on sex, population, race, tissue type, tissue state, and disease subtype (31–34).

Several lines of evidence, both computational and experimental, support the functional importance of isomiRs. Intuitively, this is not surprising considering that isomiR profiles provide a richer and more granular representation of the molecules produced from each miRNA locus compared to the single molecule, the “archetype,” that one finds listed in public databases. As we exemplified in the case of breast cancer, isomiRs are more suited to capture breast cancer heterogeneity than the archetype miRNAs (35). Recently, others and we also showed that distinct isomiRs originating from the same miRNA arm can target multiple distinct genes and molecular pathways (31,34,35).

The Cancer Genome Atlas (TCGA) initiative has been successfully integrating miRNA profiles with messenger RNA (mRNA) expression and genome-wide sequence information to further explain disease subtypes. More than 11,000 samples from 32 cancer types have been profiled today at the levels of miRNA, mRNA, protein, epigenome, etc. Tools were developed to analyze TCGA’s RNA-seq datasets and have generated what is a unique and rich resource for this kind of research (36).

Our recent analyses of expression profiles from hundreds of individuals showed that “how many” and “which” isomiRs are produced from a given miRNA locus depends on the locus and the tissue, among other variables (31,35). This is in agreement with previous reports that the archetype miRNAs of miRBase are specifically expressed in some tissues but absent from other (30,37). These findings by others and us suggest that miRNA expression signatures are complex and dynamic. Additionally, these findings support the possible use of isomiRs as biomarkers and prompted us to investigate their use as such. The ideal isomiRs to use as features in a biomarker signature should be present *primarily* in the cancer being studied and *largely* absent from the other cancer types.

Here we describe our findings from a pan-cancer analysis of TCGA’s short RNA-seq datasets. Specifically, we evaluated the ability of binary profiles that we built on isomiRs to discriminate among the 32 TCGA cancer types. For protein-coding transcripts, binary profiles were shown previously to exhibit robustness to noise and to contain enough information to distinguish among tumor types (38,39) and among tissues (40). For isomiRs, we generated “binary isomiR profiles” as follows: after thresholding in an adaptive manner, we <u>ignored</u> the isomiR’s *actual* level of abundance and instead declared it *present*, if its expression exceeds threshold; otherwise, we declare it *absent*. We also evaluated the ability of “binary miRNA-arm profiles” to discriminate among the 32 TCGA cancer types. In this case, we declare either the 5p or the 3p miRNA arm *present*, if at least one of its isomiRs is *present* above threshold, and *absent* otherwise. Clearly, the miRNA-arm representation greatly reduces the information that the analysis is allowed to use.

## MATERIALS AND METHODS

### Data acquisition and correction

We quantified the TCGA isomiR expression data of 10,271 samples at the molecule/isomiR-sequence level. In order to do this, we took the publicly downloadable loci-based *isoform.quantification.txt* files from the TCGA datasets (downloaded from the TCGA data portal https://tcga-data.nci.nih.gov on August 06, 2015) and converted them to be molecule/sequence based. Importantly, our pool of candidate biomarker miRNA loci includes miRBase as well as those hairpin arms of miRBase for which we reported recently that they are expressed in various tissues (30). Prior to the analysis, we applied corrections to account for mature sequences that could originate from any of several known miRNA paralogues. We also corrected for the fact that the *isoform.quantification.txt* files made available by TCGA often list only a subset of possible loci in the case of miRNA paralogues. Importantly, even though we counted the expression of miRNA paralogues once (thereby avoiding multiple counting) we maintained the labels of all possible paralogues throughout the analysis.

For our analyses, we used all TCGA samples from 32 cancer types. We excluded from our analyses all samples that were specifically annotated as potentially problematic samples (*file_annotations.txt* files of the Clinical Data from the TCGA data portal, downloaded on 28 October, 2015), resulting in 10,271 samples. We also removed non-tumor samples, e.g. normal adjacent tissue or metastastic samples, including only samples that had a sample infix of ‘01’ or ‘03’ in the TCGA barcode name.

### Binarized isomiR and miRNA arm profile

We worked on a per-sample basis in order to generate the binarized isomiR profiles. Specifically, for each sample independently we considered the top 20% most expressed isomiRs as ‘present’. To generate the binarized miRNA profiles, we collapsed the information at the arm level: if there was at least one isomiR (originating from the respective arm) that was characterized as ‘present’, then the arm was also marked as ‘present’. For cases in which one isomiR could be mapped to more than one miRNA arms, we further merged them into meta-arms, i.e. collections of arms that were sharing all their common isomiRs.

### Statistical and machine learning analyses

Analyses were done in R and Python. Specifically, hamming distance was calculated with the *hamming.distance* function of the *e1071* package, while all other distance metrics of hierarchical clustering (HCL) were performed with the *hcluster* function of the *amap* package. Visualization of dendrograms was performed with the *dendextend* package of R. *X*^2^ tests were performed and P values were corrected to FDR values. Binarized profiles significance using *X*^2^ tests was further filtered so that the absolute difference between the percentage (%) of samples containing the isomiR or miRNA arm in one cancer, but not the other, had to be greater than or equal to 80%. Networks were visualized using the *igraph* package in R.

Support Vector Machines (SVMs) were run with the *svm* function of the *e1071* package in R with linear kernel function and with allowed probability predictions. After the SVM model was trained, the probability vectors (one per sample) were computed for each sample in the test set. Each vector has 32 elements each one representing the probability that the given sample is of the respective cancer type. The sample is classified to the cancer type with the highest probability. If the probability < 0.5 for all cancers, then we assign the sample in the ‘Other’ category. Sensitivity and FDR scores were estimated separately for each cancer and separately for each iteration. Sensitivity was defined as the number of true positive classifications divided by the total number of samples, while FDR was calculated as the number of false positive samples divided by the number of samples identified as cancer, i.e. non-Other. For the histograms of Figs. 3C, 3D, 5C and 5D, we averaged the sensitivity and FDR scores per iteration. For Supp. Fig. S4, we correlated (Spearman’s *rho* coefficient) the sensitivity and FDR with the number of samples in each cancer and plotted the distributions as a histogram. The VI scores were computed separately for each isomiR or separately for each miRNA arm as the average of the squared values of the weights across all pairwise SVM comparisons and then were scaled to 1 by dividing by the maximum score. RandomForest was run with the H2O package in R.

PubMed entries were identified per miRNA gene. The unique gene identifiers in the Gene database of NCBI were retrieved and the number of links to PubMed entries was counted (current as of October 07, 2016).

## RESULTS

### Preliminary material and definitions

We analyzed the isomiR expression profiles for 10,271 samples from 32 cancer types (see Methods for details). To deal with miRNAs with multiple genomic copies (paralogues), we worked at the level of the sequenced reads: thusly, for isomiRs whose sequences exist at more than one genomic locus we kept one representative instance avoiding multiple counting. Consequently, we represented each sample using an expression vector with as many dimensions as the number of distinct isomiR sequences that are expressed in the sample.

As mentioned at the end of the Introduction, we intentionally focus on binary isomiR profiles, i.e. profiles that simply list an isomiR as present or absent. We determined an isomiR’s presence or absence independently for each sample and without any influences by the isomiR’s genomic origin. Binarization of isomiR abundances proceeded as follows: within the sample at hand, we considered as “present” the top 20% most abundant isomiRs; all other isomiRs were labeled “absent.” Drawing the line at the top 20%, represented an average threshold of ~10 reads per million (RPM), i.e. 10 RPM, which is a stringent threshold (Supp. Fig. S1).

In addition to working with the binarized profiles of isomiRs, we also explored an alternative scheme, namely “miRNA-arm binarization.” This representation scheme collapses the information captured by multiple isomiRs into a single statement of “present” or “absent.” Specifically, if a miRNA arm, either 5p or 3p, had at least one of its isomiRs labeled “present,” then this arm was also labeled as “present,” otherwise it was labeled “absent.”

### Statistics of binarized isomiRs

By processing the 10,271 normal and tumor TCGA samples, we accumulated a total of 7,466 isomiRs that passed threshold in at least one sample. These isomiRs arise from 807 arms that correspond to 767 miRNA loci from miRBase and 40 novel miRNA genes as we reported in Londin *et al*. (30). Supp. Tables S1 and S2 list the binary expression profiles for isomiRs and miRNA arms, respectively. Our analyses were carried out on the samples corresponding to primary solid tumors (sample infix ‘01’ in the TCGA sample barcode) except for Acute Myeloid Leukemia (LAML) where blood-derived samples were used (sample infix ‘03’).

By analyzing the 7,466 isomiRs that occupy the 20% most abundant positions in at least one sample, we found that the vast majority of them (90.2%) are present in fewer than half of the analyzed tumor samples. Only 48 out of the 7,466 isomiRs are present in all datasets (Supp. Fig. S2A). Interestingly, 11 of the 48 isomiRs arise from loci that belong to the let-7 family of miRNAs. Other isomiRs that are present in many of the analyzed datasets arise from widely-studied miRNA loci including miR-21, miR-29, miR-30, the miR-17/92 cluster and its paralogues. For individual miRNA loci, the distribution of their isomiRs varied greatly across samples. For example, let-7 isomiRs were “dichotomized:” one subset is present in most of the TCGA datasets whereas a second subset is present in fewer than 25% of the datasets (Supp. Fig. S2B).

A significant portion (58.8%) of the 7,466 isomiRs is present in fewer than 100 samples each (Supp. Fig. S2A). Moreover, 77.5% of the 7,466 isomiRs are present in at least two of the 32 distinct cancer types (Supp. Fig. S2C). These findings suggest that the expression of many of the identified isomiRs has a cancer-specific dimension.

Fig. 1A depicts as a heatmap the variation in the number of isomiRs from a given locus. Only the top 70 loci from the standpoint of the isomiRs they produce are shown in the heatmap. As can be seen, let-7a-5p consistently produces numerous isomiRs independent of cancer type. MiR-21-5p and miR-30a-3p also produce many isomiRs in many of the 32 analyzed cancers. Ovarian cancer (OV) in the case of miR-21-5p, and acute myeloid leukemia (LAML) in the case of miR-30a-3p are notable exceptions to this observation. Analogously, we also observed several miRNA arms that produce numerous isomiRs in some cancers only. The 5p and 3p arms of miR-9 are a characteristic such example: both arms produce numerous isomiRs in lower glade glioma (LGG).

**Fig. 1.**
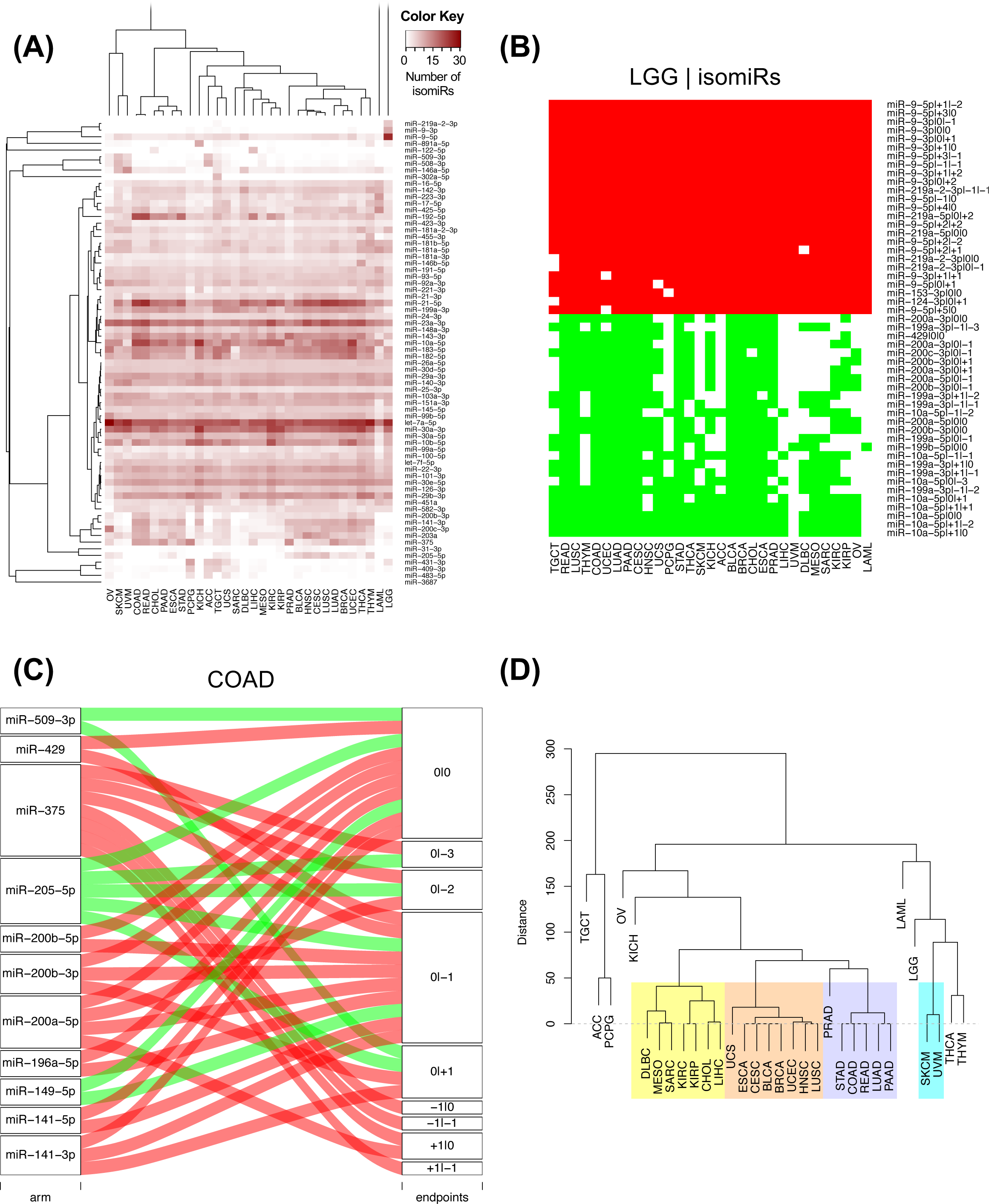
Differentially present isomiRs among different cancer types. (A) Heatmap of the number of isomiRs per miRNA arm. The darker the color of each cell, the higher the number of isomiRs that the respective arm (rows of the heatmap) produces in the respective cancer type (columns of the heatmap). (B) Differential presence of isomiRs in lower grade glioma (LGG) tumor samples as compared to the rest of cancer types. Red indicates that the isomiR (row) was found as ‘present’ in LGG as compared to the respective cancer type (column), while green indicates ‘absence’ in LGG. The data from all possible pairwise comparisons is included in Supp. Table S3. (C) Overlap between miRNA arms and isomiRs in the comparison of colon adenocarcinoma (COAD) with the rest of the cancer types. Red indicates that both the isomiR and arm were ‘present’ in COAD as compared to at least one other cancer type, green indicates they were ‘absent’. For example, miR-205-5p was ‘absent’ in COAD as well as its five isomiRs, miR-205-5p|0|0, miR-205-5p|0|-3, miR-205-5p|0|-2, miR-205-5p|0|-1 and miR-205-5p|0|+1. (D) Hierarchical clustering (complete method) considering the number of differentially present isomiRs as the distance between cancer types. Colored clusters are described in the main text.

### IsomiR production is cancer-dependent

We studied systematically the binary differences of presence/absence of abundant isomiRs among cancers by conducting all possible pairwise comparisons among 32 cancers. For each comparison, we performed *x*^2^ tests, suitable for comparison of binary data, for all isomiRs in the two given cancers. To focus on the most discriminatory isomiRs, we imposed a False Discovery Rate (FDR) threshold of 0.1% and further required that the percentage of samples in each cancer that contain the isomiR differ by at least 80%. We were able to identify several isomiRs that were significantly present in one cancer and absent from many of the remaining ones (Supp. Table S3). Fig. 1B illustrates this observation using LGG as an example. As already mentioned above (Fig. 1A), isomiRs from the miR-9-3p arm are *present* in LGG samples and *absent* from nearly all other cancers. The opposite holds true for isomiRs from the miR-10a-5p arm and the miR-200 family: they are *absent* from LGG and *present* in 71-93% of the other cancers. Another characteristic example can be seen in Supp. Table S3: in testicular germ cell tumors (TGCT), the miR-302 family and miR-371/372/373 cluster express several isomiRs absent from nearly all other cancers.

### MiRNA-arm transcription is cancer-dependent

Noticing the co-presence/co-absence of isomiRs from the same miRNA arm in some cancers, we tested the hypothesis that the miRNA arms themselves are also differentially present among cancer types. We repeated the previous *χ*^2^ analysis for the binarized profiles of miRNA arms (see above for definitions) and were able to largely replicate the results that we obtained at the isomiR level (Supp. Table S4). Colon adenocarcinoma (COAD) provides a characteristic example. At the isomiR level, several isomiRs from the miR-215-5p arm were found to be COAD-specific when compared to the other cancer types (Supp. Table S3). Looking at miRNA arms only, we find that the 5p arm of miR-215 also exhibits the same trend, i.e. its production of isomiRs is specific to COAD (Fig. 1C and Supp. Tables S3 and S4).

### Binarized isomiR profiles can be used to hierarchically cluster cancers

First, we examined how well we can classify the 32 cancer types using binarized isomiR profiles and hierarchical clustering (HCL). As a distance metric between two cancers, we used the Hamming distance between the respective binary isomiR profiles (Supp. Table S3). Essentially this measures the isomiR differences (present → absent, absent → present) between the two cancers being compared. The resulting dendrogram is shown in Fig. 1D. In it we observe several interesting clusters. A first cluster (light purple background) contains almost all the adenocarcinomas, like pancreatic ductal (PAAD) and prostate adenocarcinoma (PRAD). A second cluster (light orange background) encloses the breast (BRCA) and bladder (BLCA) cancer types along with the squamous cell carcinoma of lung (LUSC) and head and neck (HNSC). A third cluster (light yellow background) includes cancers of the kidneys (renal clear cell carcinoma, KIRC; renal papillary cell carcinoma, KIRP), liver (hepatocellular carcinoma, LIHC) and the bile duct (cholangiocarcinoma, CHOL). We also note the clustering of the uveal (UVM) and skin (SKCM) melanomas (cyan background).

The above clustering into tumor groups implies a small Hamming distance and indicates similarities in the profiles of abundant isomiRs. By extension, the profile similarities imply commonalities in the respective molecular physiologies. However, this univariate analysis is not suitable for tackling the multidimensional question of cancer classification.

### Conventional multivariate clustering cannot separate all cancer types

We next embark on multivariate statistical approaches to evaluate the hypothesis that the binarized isomiR and binarized miRNA-arm abundance profiles can be used for tumor discrimination and classification. After computing Hamming distances between pairs of samples (not pairs of cancers) using the respective binarized isomiR profiles, we carried out HCL. We were able to discriminate up to seven cancers using the binarized isomiR profiles (Fig. 2A). Collapsing isomiR profiles into binarized miRNA-arm profiles did not provide any additional discriminatory ability (Fig. 2B).

**Fig. 2.**
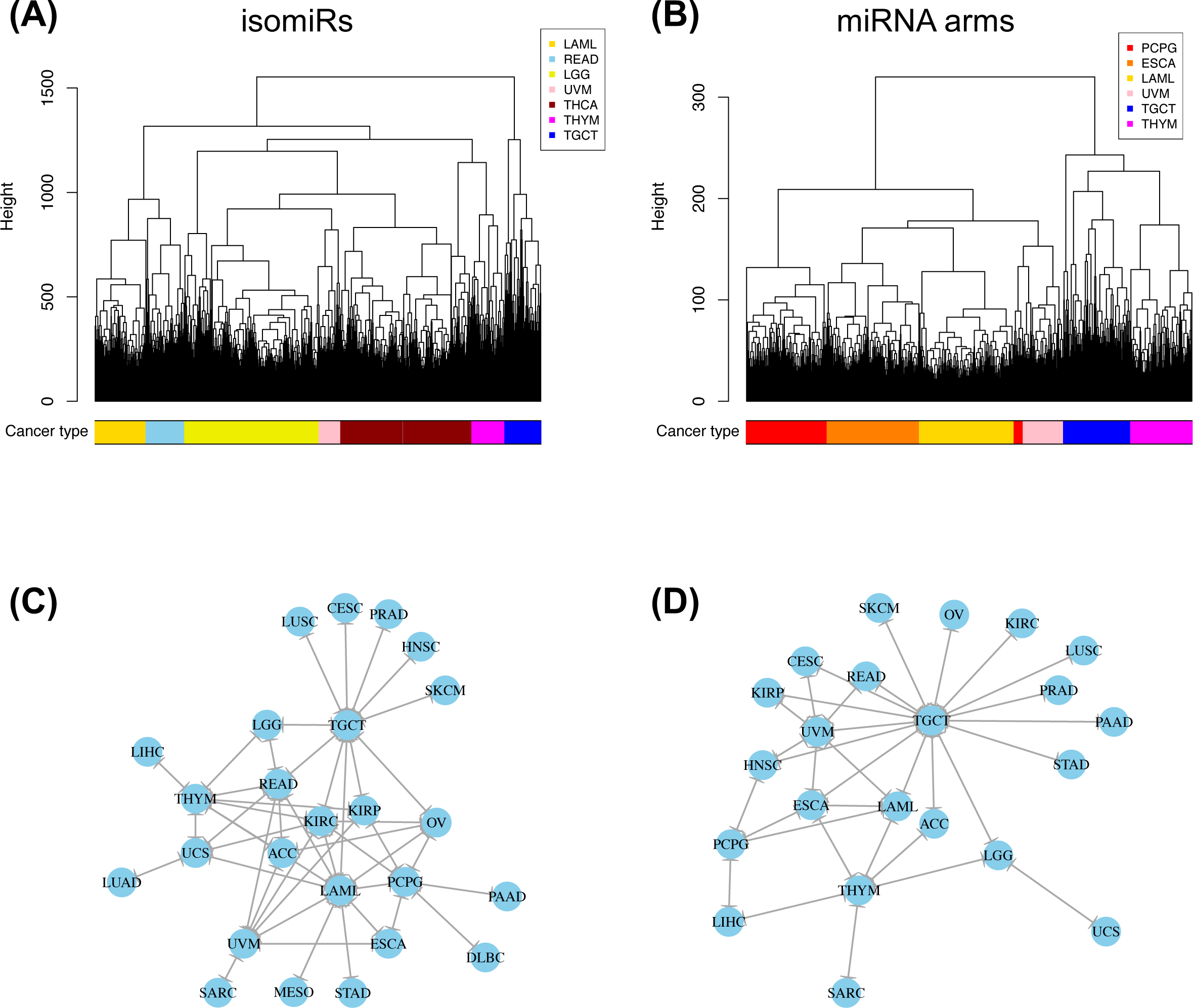
Multivariate hierarchical clustering on the binary expression vectors. (A and B) Hierarchical clustering (hamming distance as metric) on the isomiR (A) or miRNA arm (B) profile of samples from different cancer types. The leaves of the dendrogram are tumor samples. The colored bar indicates the cancer type of the respective sample. (C and D) Networks of all potential pairwise discriminations using hierarchical clustering (hamming distance as metric) on the isomiR (C) or miRNA arm profile (D). Two nodes (cancers) are connected if and only if the corresponding samples were found to form two separate clusters, with the samples of one cancer clustered distinctly from the other.

To investigate the upper limit of using HCL and Hamming distances at the sample level, we considered all possible cancer pairs and performed all comparisons among the respective samples. In each case, we examined whether each cancer’s samples would form their own cluster. Figs. 2C (isomiRs) and 2D (miRNA arms) illustrate the outcome of this analysis. In the shown networks, each node is a cancer type. Two nodes are linked with an edge if and only if the corresponding cancers can be distinguished from one another (large Hamming distance between samples from the respective cancers). LAML, TGCT and thymoma (THYM) appear as central hubs in these networks: this means that these cancers can be distinguished with relative ease from several other cancer types. We note the absence of nodes for e.g. colon adenocarcinoma (COAD) and thyroid cancer (THCA) from these networks. This indicates that, using the current clustering model (Hamming distance + HCL), COAD and THCA cannot be distinguished from other cancers. Also, not surprisingly, binarized isomiR profiles (Fig. 2C) can separate several more cancers from one another compared to binarized miRNA-arm profiles (Fig. 2D).

### Binarized isomiR profiles can discriminate amongst cancers

Support Vector Machines (SVMs) have been gaining popularity, due to their capacity for multi-class classification in many different contexts (41–45). SVMs are intrinsically designed for binary classifications, i.e. for finding the best hyperplane that separates <u>two</u> *a priori* defined clusters. For our multi-cancer classification, we used an approach analogous to PhyloPythia, our previously published method for classifying metagenomes (41,42). In particular, we build 496 SVMs for each of all possible “cancer-type-X vs. cancer-type-Y” pairwise comparisons, and then integrate the information into a single model by attaching probabilities to the classification outcome.

We split our 9,293 tumor datasets into training sets (used to *construct* each model) and test sets (used to *evaluate* each model), as is general practice in machine learning. Specifically, for each cancer type in turn, we formed a training set that comprised 60% of the type’s samples. The remaining 40% of the sample’s for the cancer type at hand formed the test set. For each cancer type, we used the respective “training” samples to build an SVM aimed at separating the cancer type at hand from the remaining 30 cancer types.

We built 496 SVM models using the “binarized isomiR profiles” and another 496 models using the “binarized miRNA-arm profiles.” The isomiR SVM models were evaluated separately from the miRNA-arm SVM models. For the 496 SVM models being considered, we presented each test sample to each of the 496 SVMs in turn and used their output to build a 32-dimensional vector of probabilities: the *i*-th element of the vector is the probability that the test sample at hand belongs to the *i*-th cancer type. We imposed a probability threshold of 0.5: if the *i*-th element of the probability vector was ≥ 0.5 then the test sample was classified as belonging to the *i*-th cancer type. If none of the 32 probabilities reached the 0.5 level, then the test sample was assigned to the ‘Other’ category. The ‘Other’ category contains false negatives as well as samples that truly do not belong to the 32 considered cancer types.

We repeated each cycle of SVM model building and testing 1,000 times. In each iteration, we used a different set of randomly selected samples for training. Fig. 3A shows a heatmap that summarizes the average prediction performance of the SVMs that are based on the binarized isomiR profiles. Each row designates the cancer type to which the test sample belongs and each column designates the predicted cancer type. A perfectly-specific classifier should not generate any non-diagonal entries. A perfectly-sensitive classifier should not generate any entries in the “Other” category. As can be seen, the binarized isomiR profiles can clearly discriminate among cancer types and to correctly classify each sample to its correct origin. There is one notable instance of seemingly diminished performance that involves several rectum adenocarcinoma (READ) tumors that were misclassified as COAD tumors. In reality, this is not unexpected considering that READ and COAD tumors are molecularly similar and their distinction is largely driven by anatomy (46). Supp. Table S5 contains an example of the probability vectors for the test samples as well as the confusion matrix from one of the 1,000 iterations. It is evident that the probabilities for the correctly assigned samples were considerably high, most of them being ≥ 0.9. We also note that the 1,000 iterations built SVM models that were fairly similar to one another (Supp. Fig. S3A and S3B), indicating high stability in the potential presence of extreme outlier samples.

**Fig. 3.**
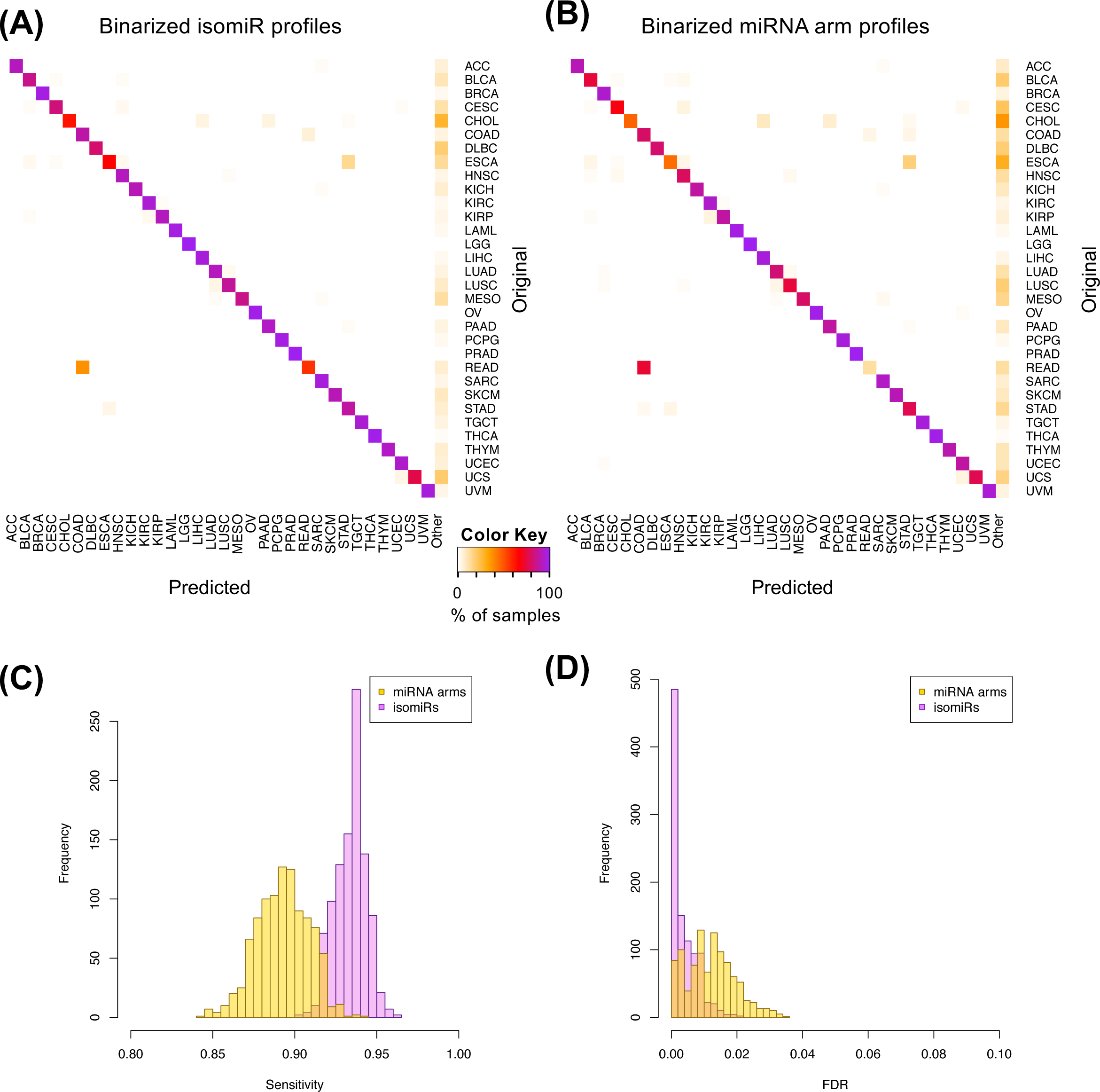
Support Vector Machines correctly classify 32 cancer types. SVM classification using the binarized isomiR (A) or the miRNA arm (B) expression profile. Each row of the heatmap represents the original and each column the predicted cancer class. The color of each cell in the heatmap is proportional to the percentage (%) of samples originally as the cancer type in the respective row to be predicted as the cancer type of the respective column. The % is calculated as the average across 1,000 iterations. (C-D) Sensitivity (C) and FDR (D) scores for the SVM models built using the binarized isomiR (magenta) or miRNA arm (yellow) expression profiles.

Fig. 3B shows an analogous heatmap that summarizes the average prediction performance of the SVMs that are built using the binarized miRNA-arm profiles. As can be seen, the classification remains largely successful. This suggests that simply interrogating whether a miRNA arm produces isomiRs can provide good performance when classifying cancer samples.

Finally, we quantified the performance of the 1,000 SVM models by building the distribution of the respective sensitivity and False Discovery Rate (FDR) scores. We did this separately for the binarized isomiR and miRNA-arm SVM models. Fig. 3C shows the resulting distributions for sensitivity and Fig. 3D for FDR. It is evident that binarized isomiR profiles are considerably more effective in correctly classifying tumor samples showing a mean sensitivity score of 93%. Even though lower (87%), the average sensitivity of the SVMs that were based on the binarized miRNA-arm profiles is fairly high in absolute terms. Indeed, one should consider here the magnitude of the task and how little information is actually used in this case when building these SVM models. The FDR scores also supported the effectiveness of the SVM-based classification. Specifically, the “binarized isomiR profiles” exhibited a mean FDR of 3% whereas the “binarized miRNA arm profiles” exhibited a mean FDR of 5%.

### Validating the resulting SVM classification

To ensure that the achieved SVM classification is not artificial, we carried out two tests. In the first test, we kept the number of ‘present’ isomiRs constant but randomly rearranged them in each sample. We then proceeded with building our 1,000 isomiR-based SVM models and tested them with the “correct” test samples. As expected, all of the test samples, in all 1,000 iterations, were assigned to the ‘Other’ category (Supp. Fig. S3C).

In the second test, we shuffled the labels of the training samples prior to training each SVM model with the binary isomiR profile. As before, we built 1,000 SVM models and tested them with the “correct” test samples. Doing so resulted in the complete collapse of the model, consistent with our first step (Supp. Fig. S3D).

We also note that there is a weak correlation between the observed sensitivity and FDR scores with the number of samples in each cancer type (Supp. Fig. S4), indicating that the success or failure of classification is not driven by the uneven sample sizes.

Based on the outcome of these two tests, we conclude that the classification results depicted in Fig. 3 are not driven by random or accidental events. Instead, the binarized isomiR and miRNA-arm profiles appear to carry actionable information.

### The most discriminatory isomiRs and miRNA arms are <u>not</u> among those frequently-studied

As the SVM attempts to identify the best-separating hyperplane in the multi-dimensional space, some of the features are given more weight than others. In our case, these features would be tantamount to specific isomiRs and specific miRNA arms respectively.

To identify those isomiRs that were deemed most significant in separating the various cancer types, we ran the SVM method using as training set the whole TCGA dataset and extracted the variable importance (VI) score for each variable as the mean of the squares of the feature weights (47) of all pair-wise comparisons (Supp. Table S6). We repeated the same analysis for miRNA arms and identified those with the highest VI values (Supp. Table S7).

Among the isomiRs, we found that two isomiRs from the 5p arm of miR-205 were deemed most important by the isomiR-based SVM classifiers (Supp. Table S6). These were followed by several isomiRs from both arms of miR-141. Notably, we observed a trend for agreement between the SVM models built on isomiR profiles and miRNA-arm profiles respectively with regard to the miRNA loci that the two models deem important (Supp. Tables S6 and S7). The loci include miR-205, miR-141, and miR-200c.

To validate these findings, we used the RandomForest algorithm, which has been shown able to identify significant variables for classification (48). We found the VI scores from RandomForest to be strongly and positively correlated with the VI scores from the SVM models (Supp. Table S6): Spearman rho correlation coefficient 0.886 (P-val<0.01). The correlation improves further to 0.932 (P-val<0.01) when we compare the VI scores obtained from the binarized miRNA-arm models (Supp. Table S7). The fact that a second independent algorithm validates the SVM conclusions adds further support to the relevance of using binarized profiles.

Having confirmed with two independent machine-learning tools the VI scores, we associated the corresponding molecules with the number of PubMed. For this step, we specifically used those miRNA loci that have entries in the Gene database of NCBI and retrieved the number of PubMed entries associated with each miRNA gene (see Methods). Fig. 4A shows the result of this association for SVMs whereas Fig. 4B shows the analogous results for RandomForest. We observed a mean of 52 publications per isomiR and 30 publications per miRNA arm. A striking result of this analysis is that both arms of the mir-21 precursor are each associated with the highest number (689) of publications. However, with regard to their ability to classify cancer samples, both the SVM and RandomForest models assign them a considerably low VI score (little discriminatory power). It is also important to underline that both the SVMs and the RandomForest deemed miR-944 to be among the most important miRNAs for cancer classification: miR-944 currently has only six PubMed entries (Supp. Table S7). Other examples of discriminatory miRNAs with few PubMed entries include miR-429 (87 entries), miR-192 (46 entries), miR-194 (10 entries), and miR-135b (40 entries). Lastly, we note a similarly weak correlation between the number of PubMed entries and the number of times the miRNA arm is found differentially present between two cancer types (Fig. 4C).

**Fig. 4.**
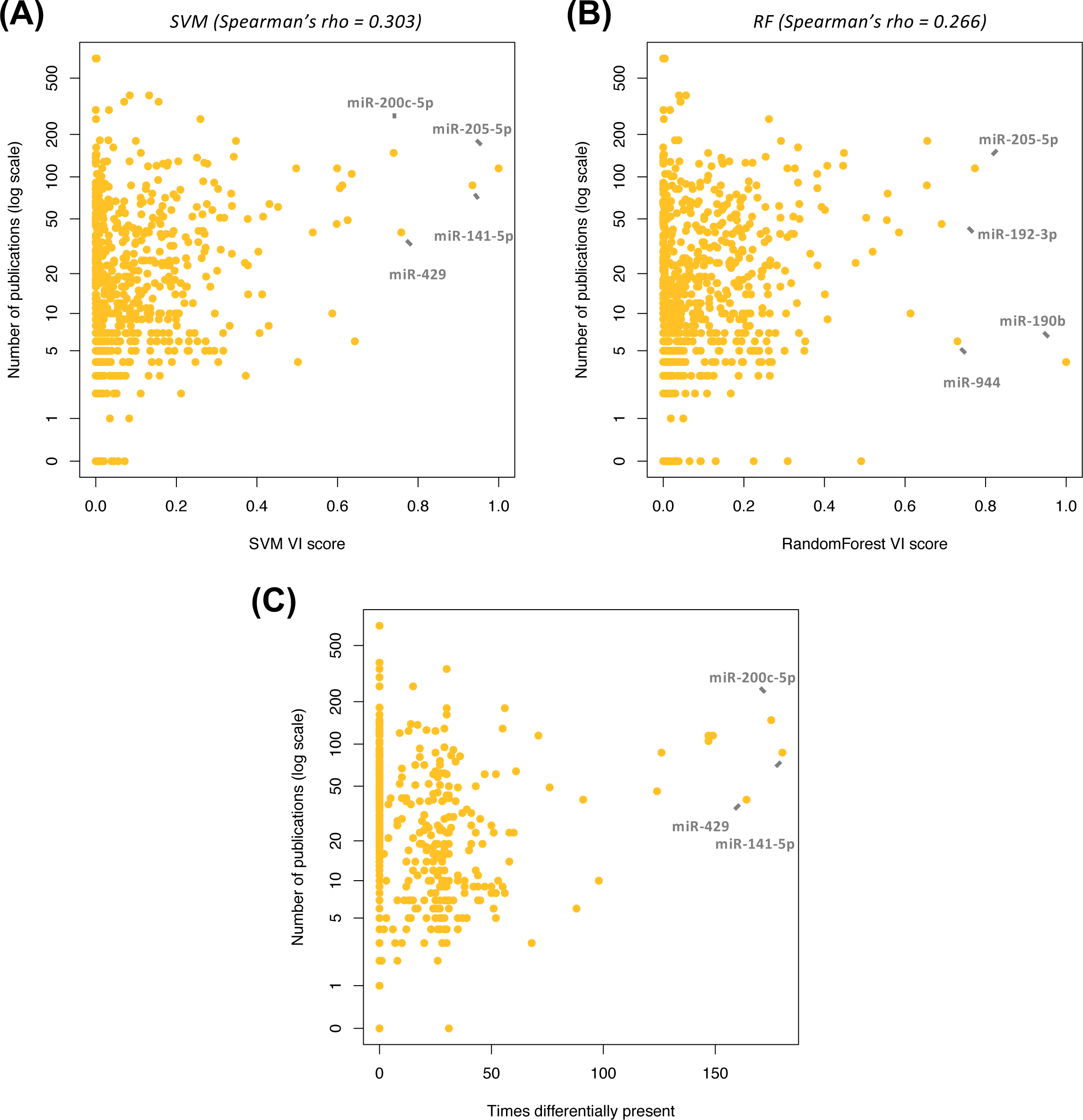
Number of publications does not correlate with importance in classification. Number of publications against the variable importance (VI) score as calculated in the SVM (A) or RandomForest (B) classification model based on miRNA arms. Spearman correlation coefficients: for SVM: 0.303, for RandomForest: 0.266. Number of publications against the times each miRNA arm was found differentially present (Supp. Table S4). Spearman correlation coefficient: 0.215.

Lastly, we repeated the above analysis with one change. Specifically, we examined the correlation of the VI score with the number of times that the isomiR, or miRNA arm respectively, is found to differentially present in a pairwise comparison. We found that those of the isomiRs and miRNA arms that had the most impact on cancer classification were the ones found to be differentially present among many cancer types (Supp. Figs. S5A and S5B). I.e. those isomiRs and miRNA arms that are uniquely present or absent in one cancer type are <u>*not*</u> the most impactful ones. We obtained similar results for both isomiRs and miRNA arms under the RandomForest model as well (Supp. Figs. S5C and S5D).

Summarily, the above findings suggest that the current body of literature includes a limited number of studies of miRNAs that, as per our analysis, have the most potential to serve as cancer-specific biomarkers.

### Use of a reduced set of features preserves the ability to classify with binarized profiles

The above-mentioned SVM and RandomForest models considered all 7,466 present isomiRs (isomiR profiles) or all 807 present miRNA arms (miRNA-arm profiles). As we discussed in the previous section, only a relatively small number of isomiRs and miRNA-arms are of considerable value in cancer classification. Based on these observations, we investigated the possibility that we could obtain reasonable classification results using a reduced set comprising the most important features (isomiRs or miRNA arms, Fig. 5A and 5B, respectively).

**Fig. 5.**
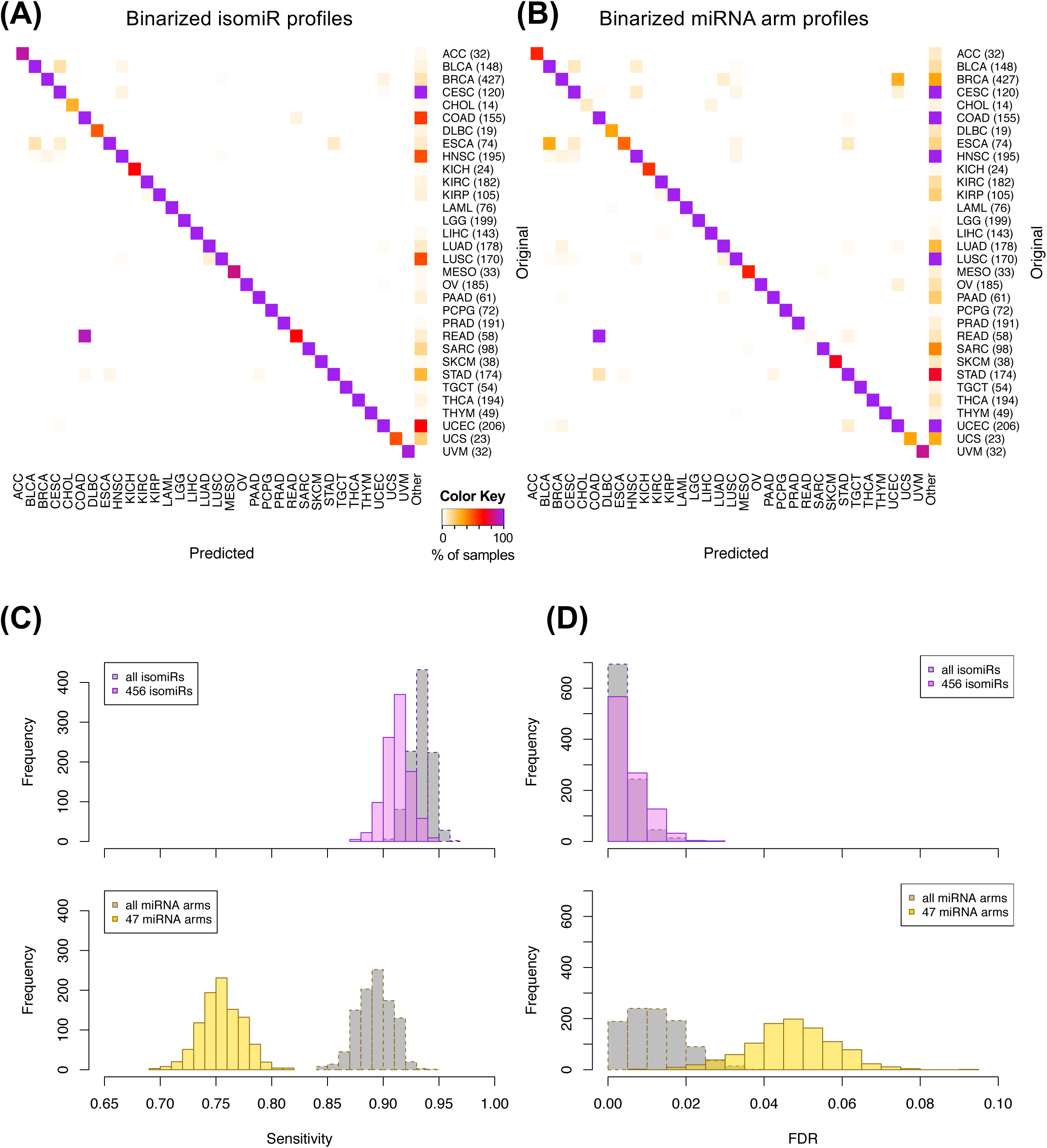
Support Vector Machines classification with a reduced list of isomiRs is more robust than one of miRNA arms. SVM classification using the reduced list of 456 isomiRs (A) or 47 miRNA arms (B). Each row of the heatmap represents the original and each column the predicted cancer class. The color of each cell in the heatmap is proportional to the percentage (%) of samples originally as the cancer type in the respective row to be predicted as the cancer type of the respective column. The % is calculated as the average across 1,000 iterations. The numbers in parenthesis show the number of test samples in each cancer type. (C-D) Sensitivity (C) and FDR (D) scores for the SVM models built using the reduced binarized isomiR (top graph) or miRNA arm (bottom graph) expression profiles. The distributions with the full isomiR and miRNA profiles are shown in gray.

We intersected the top 10% most important isomiRs of the SVM model with the top 10% isomiRs of the RandomForest and obtained 456 isomiRs (Supp. Table S8). Using these 456 features (instead of the original 7,466) we repeated the multi-cancer SVM-based training and classification. We found that even with this reduced set of isomiR features we maintained our ability to correctly classify samples (Fig. 5A) – the concomitant sensitivity following 1,000 training/testing iterations decreased by a mere 3% to 90% (Fig. 5C) and the FDR by a mere of 1% to 4% (Fig. 5D). Much of the error originated from the higher uncertainty of the method to classify samples, as evidenced by the higher number of samples clustered as “Other” (Fig. 5A). It is important to stress here that the diagonal of the heatmap shown in Fig. 5A attracts the vast majority of the tested samples (correct classifications) and supports the use of isomiR-profile-based SVMs in cancer classification.

We also repeated the analysis using a reduced set of most important miRNA-arm features to build our SVM models. We intersected the top 10% most important miRNA arms of the SVM model with the top 10% miRNA arms of the RandomForest and obtained 47 arms (Supp. Table S8). This signature of miRNA arms was sufficient to classify the samples to their respective cancer type (Fig. 5B) at the expense of a modest penalty in sensitivity (decreased to 78% from 87% when all features were used) and FDR (increased to 8% from 5% when all features were used) as seen in Figs. 5C and 5D, respectively. These results suggest that the reduced set of miRNA arm features retains a considerable ability to classify cancer samples; however, it does not reach the levels achieved by isomiRs. On the other hand, the miRNA-arm-based SVMs achieve these results using ten times fewer features than isomiRs.

## DISCUSSION

In this study, we examined the relevance of binary isomiR and miRNA-arm profiles in distinguishing amongst multiple cancer types. This work was spurred by previous observations that miRNA expression profiles can be tissue-specific (28,29,37,49) and tissue and cell type differences can be adequately described by only the presence or absence of RNA transcripts (39,40). We centered this work on isomiRs, miRNA isoforms whose importance we demonstrated in recent publications (31,35). Specifically, we sought to determine how well binary profiles of isomiRs, and of miRNA arms, can describe cancer types. To this end, we leveraged the TCGA datasets due to the standardized protocols and data availability (36).

A first result that emerged from our analysis was the identification of several instances of cancer-specific presence or absence of expression for isomiRs or for their corresponding hairpin arms. The most striking example is miR-9 whose isomiRs are uniquely present in LGG tumor samples. This miRNA is highly expressed in the nervous system and has evident important roles in neuronal development and diseases (50,51) supporting our findings of its tissue-specificity. Another miRNA that also exhibited similar characteristics in LGG was miR-219 and, intriguingly, this miRNA has been implicated in neural differentiation processes (52). We were also able to identify additional miRNAs that have almost unique expression in some tissues, like the liver-specific arms of miR-122 (53). Our unbiased global approach also identified potential cancer-type-specific miRNAs: examples include miR-671 for ovarian cancer, and our novel miRNA ID00737-3p for THCA (Supp. Tables S3 and S4). Ovarian cancer was also an interesting because a relatively high number of isomiRs and miRNA arms were <u>*absent*</u> compared to other cancer types. The opposite was true for TGCT tumor samples where several isomiRs were present exclusively, e.g. isomiRs from the miR-302 family (Supp. Tables S3 and S4) that has important roles in stem cell pluripotency and cell reprogramming (54,55).

It is important to stress that causative links for the above observations cannot be identified based on our analysis. Also, the cancer-specific expression can not be guaranteed as the isomiRs and the miRNA arms can preserve tissue-specific expression trajectories, even in the cancerous state. This is not unexpected considering that cellular context matters in cancer biology (56) and could be contributing to cancer-type differences. As the TCGA projects were largely focused on tumor classifications, limited normal samples were collected by the various consortia. Further studies with adequate numbers of normal samples will be needed to decouple the “normal” from the “cancer” signal, similarly to what was done previously (28).

From a biomarker perspective, tissue specificity is of great importance, as miRNAs that are ubiquitously expressed in multiple tissues are not appropriate candidates for this role. In this regards, our work represents a first and much-needed step towards a global assessment of the usefulness of miRNAs and their isoforms as biomarkers. Two characteristic examples here are the mir-21 locus – the miR-21-5p miRNA has the highest number of publications, and the let-7 family several members of which are present in all the samples and all the cancer types of our study. These observations suggest that these miRNAs are not adequately specific to be biomarkers (57). Such complications led several groups to suggest the use of “miRNA panels” as biomarkers rather than using one or a handful of miRNA molecules (58–60).

Having established that qualitative differences among cancer types exist and are meaningful, we employed multivariate and machine learning tools to build classification models and predict the cancer type solely based on binary expression information of isomiRs or miRNA arms. We found that building SVMs from the isomiR binary profile were capable of correctly predicting the cancer class with more than 90% sensitivity with an FDR < 5% (Fig. 3). SVMs built from isomiRs outperformed SVMs built from miRNA-arm profiles (Fig. 3), even when we reduced the number of isomiR features by one order of magnitude (Fig. 5). When we focused only on the most important miRNA arms, we observed lower prediction rates, which suggests that a proportionally large number of miRNA arms is necessary for multi-cancer classification purposes. Independent evidence that isomiRs have higher discriminatory power that miRNA arms is currently limited. In a recent work, Koppers-Lalic *et al*. found that miRNA isoforms are able to improve the specificity in prostate cancer detection, including isomiRs from the miR-204-5p, miR-21-5p and miR-375-3p arms (61). In addition, our previous work in breast cancer showed that isomiRs can better distinguish normal breast tissue from BRCA tumors by comparison with archetype miRNAs (35). A potential explanation for the higher predictive value of isomiRs can be that quantification and modeling at the miRNA arm level inherently discards information, which can be rooted in biological mechanisms that may specifically affect isomiR levels (62–64).

A potentially significant contribution of the current work can be in the field of cancers of unknown primary site (CUP) (65–68). MiRNA profiling has evidently allowed for the identification of the primary sites of metastatic cases (69). Screening for the presence or absence of isomiRs or miRNA arms that were found significant in our analyses could further enchance the prediction accuracy.

In summary, our paradigm of binary expression signature of isomiRs further extends the current literature on cancer classifications that are based on small ncRNAs. Compared to earlier work in the field, one notable difference is that previous studies relied on continuous multi-valued expression signatures to classify samples (27–29) whereas we use a binary input. Another difference is that our models were trained with a higher number of samples and can discriminate amongst a larger number of cancer types. We also score the “importance” of each isomiR or miRNA arm vis-à-vis its contribution to the multi-cancer classification, which we calculate from two independent models. Finally, we provide a framework for cancer-type-specific isomiR and miRNA signatures that can be readily utilized for specific biomarker discovery and also for hypothesis generation on the tissue-specificity of this class of small ncRNAs.

## Author Contributions

IR conceived the study. AGT and IR designed and supervised the study with contributions from IC and EL. AGT, RM and PL downloaded and corrected the expression and clinical data. AGT performed the analyses with contributions from RM and PL. AGT and IR wrote the manuscript. All authors read and approved the final manuscript.

## Acknowledgements

The project was supported in part by a William Keck Foundation grant (IR) and by Institutional Funds.

